# The acquisition of *rmpADC* can increase virulence of classical *Klebsiella pneumoniae* in the absence of other hypervirulence-associated genes

**DOI:** 10.1101/2025.09.20.677538

**Authors:** Stephen M. Salisbury, Taryn A. Miner, Leslie A. Kent, Margaret M.C. Lam, Kathryn E. Holt, Virginia L. Miller, Kimberly A. Walker

## Abstract

*Klebsiella pneumoniae* is one of the most common causes of nosocomial infections, and the rise of drug-resistant *K. pneumoniae* strains is complicating treatment and contributing to a mounting global health crisis. *K. pneumoniae* has two pathotypes, classical (cKp) and hypervirulent (hvKp). CKp typically cause opportunistic infections in immunocompromised individuals in healthcare settings and often are multi-drug resistant. HvKp can be community acquired and cause high-mortality infections in immunocompetent individuals. Concerningly, antibiotic-resistant cKp strains with hypervirulence-associated genes and traits have recently emerged. Determining if and how hv-associated genes contribute to increased virulence of cKp strains is essential to addressing this growing threat. The *rmpADC* operon is an hv-associated locus that confers hypermucoviscosity (HMV), a key virulence phenotype, and *rmp* genes are often found in convergent strains. In this study, we aimed to determine if the *rmp* genes alone could increase the virulence of classical strains in the absence of other hv-associated genes. We introduced genetically distinct *rmp* loci from different lineages into a broad array of cKp isolates and found that, while many isolates became HMV positive, only a subset of these strains showed an increase in virulence in a mouse model of pneumonia. Sequence type and capsule type were not predictive of how *rmp* acquisition impacted the clinical isolates. Our results indicate that HMV is likely necessary but not sufficient for hypervirulence and that *rmp* sequence can influence virulence potential in classical strains.

**IMPORTANCE:** *Klebsiella pneumoniae* is a global pathogen, and gene exchange between hypervirulent (hvKp) and classical (cKp) strains is a rising threat. It is essential to understand how hvKp genes impact virulence phenotypes and identify the cKp strain backgrounds most amenable to enhanced virulence. Hypermucoviscosity (HMV) is a critical virulence factor in hypervirulent *K. pneumoniae*, conferred by the *rmpADC* locus. The *rmp* genes are encoded on mobile genetic elements and have been detected in convergent antibiotic-resistant *K. pneumoniae* strains of concern. In this study, we explored the impact of *rmp* acquisition in a broad set of classical clinical isolates. We observed that HMV appears necessary, but not sufficient, for increased virulence. Sequence type, capsule type, and HMV capacity could not predict which classical isolates gain an *rmp-*dependent colonization benefit. These insights increase our understanding of the distinctions between cKp and hvKp and further our ability to identify and treat new strains of concern.

## INTRODUCTION

*Klebsiella pneumoniae* is one of the most common causes of hospital-acquired pneumonia and a global pathogen of growing concern. *K. pneumoniae* can cause a wide variety of diseases, including bacteremia, meningitis, urinary tract infections, and liver abscesses, and it is a major cause of neonatal sepsis [1, 2]. These infections are often difficult to treat because many *K. pneumoniae* strains have acquired resistance to multiple antibiotics [3–7]. *K. pneumoniae* contributes to a larger problem: the US Centers for Disease Control and Prevention has deemed the increase in β-lactam-resistant and carbapenem-resistant *Enterobacteriaceae* to be serious and urgent issues, respectively [8], and the World Health Organization included drug-resistant *K. pneumoniae* on its 2024 Bacterial Priority Pathogens List [9].

*K. pneumoniae* is broadly divided into two pathotypes, classical and hypervirulent [1, 10]. Classical strains (cKp) are often multi-drug resistant (MDR) and typically cause nosocomial infections in immunocompromised individuals. Hypervirulent strains (hvKp) are community acquired and can cause life-threatening infections in immunocompetent individuals. While these populations are distinct, both have spread globally [11, 12]. Concerningly, horizontal gene transfer between hvKp and cKp strains has given rise to convergent *K. pneumoniae* strains that are both MDR and hypervirulent [13–15]. Convergence occurs when cKp strains acquiring hypervirulence-associated mobile genetic elements or when hvKp strains acquiring resistance to antibiotics [16–18]. The ongoing threat of strain convergence highlights the need to better understand which hv-associated genes confer hypervirulent phenotypes.

*K. pneumoniae* is a highly diverse species, with a pangenome of at least thirty thousand genes [19]. *K. pneumoniae* populations move fluidly between natural and healthcare environments, and a wide variety of strains are capable of causing invasive disease [13, 20].The genetic determinants of virulence and hypervirulence are still not fully understood and the genetic diversity of this species complicates identification efforts [21]. Known *K. pneumoniae* virulence factors include lipopolysaccharide (LPS), type 1 and type 3 pili, a polysaccharide capsule, and siderophores, all of which are typically present in both classical and hypervirulent strains [1]. Classical strains universally produce enterobactin, and approximately 40% also produce yersiniabactin [22, 23]. HvKp strains are distinguished from cKp strains by producing additional siderophores, including salmochelin and aerobactin, and exhibiting hypermucoviscosity (HMV), although these traits alone do not define hvKp [24]. HMV is a capsule-dependent phenotype characterized by uniform and abundant capsule polysaccharides [25]. HMV-positive strains are typically string-test positive, although HMV is regularly assessed with the more quantitative sedimentation resistance assay [1, 26]. HMV blocks phagocytosis by macrophages, but it is unclear how it might otherwise affect virulence [27, 28].

The HMV phenotype is conferred by the *rmpADC* operon and appears essential for full virulence of hvKp strains [27, 29–32]. RmpA is a transcriptional regulator with a LuxR-like DNA binding domain that auto-regulates the *rmp* promoter [33–35]. RmpD is a small protein that interacts with capsule export machinery to confer the HMV phenotype [27]. RmpC, another regulator with a LuxR-like DNA binding domain, increases capsule production [34]. Deletion of any of the three *rmp* genes severely attenuates mouse lung colonization in a pneumonia model [27, 34]. HMV-negative *rmpD* and *rmpA* mutants both fail to establish infections in mouse lungs. The HMV-positive *rmpC* mutant is readily cleared, but at a slower pace than the HMV-negative mutants.

The *rmp* genes, along with other virulence-associated genes of known and unknown function, are present on large virulence plasmids and mobile chromosomal elements [36]. There are five distinct lineages of *rmpADC* associated with unique plasmids or chromosomal elements: *rmp1, rmp2, rmp2A, rmp3,* and *rmp4* [37]. Representative loci from all five lineages have demonstrated functional capacity in KPPR1S, a well-characterized hvKp strain harboring an *rmp3* locus in the chromosomally integrated ICE*Kp*1. Expression of *rmp* loci from all five lineages significantly increased capsule production in a KPPR1S Δ*rmp3* mutant, and all but the *rmp4* locus fully restored HMV [37].

Given the rising threat of convergent *K. pneumoniae* strains, it is important to determine which hv-associated genes are sufficient to affect the virulence of classical strains and which classical strain genetic backgrounds might pose the highest risk for problematic convergent phenotypes. The large mobile genetic elements (MGEs) that distinguish hvKp from cKp contain numerous genes, but only a few have been explicitly linked to virulence. The *rmp* locus has long been associated with hypervirulence, and a fully intact and functional *rmpADC* gene set has proven essential for full virulence in multiple mouse models [1, 27, 31, 34]. While the *rmp* genes have been identified in convergent strains [38], their role in virulence has mostly been characterized within hvKp. In this study, we sought to determine if *rmp* is sufficient to impact the virulence of classical strains. To this end, we analyzed the effects of *rmp* expression in a wide array of genetically distinct classical clinical isolates using *in vitro* assays and a murine pneumonia model. Here, we report that *rmp* acquisition alone can promote HMV, confer hypercapsule, and increase the virulence of select cKp isolates, but the impact on virulence varies by strain background. Additionally, *rmp* lineage and cKp genetic background contribute to *rmp-*dependent virulence changes in classical strains.

## RESULTS

### Ectopic *rmp* expression in classical strains confers varied HMV levels

The *rmp* genes confer two well-characterized phenotypes associated with hypervirulence, HMV and increased capsule production (hypercapsule) [27, 29, 34]. We set out to determine if the *rmp* genes are sufficient to confer these phenotypes in multiple cKp isolates, and to ascertain if there are any functional differences across *rmp* lineages. To this end, we transformed a diverse set of eighteen cKp isolates with low copy plasmids containing a representative *rmp* locus from each of the five lineages (*rmp1, rmp2, rmp2A, rmp3,* and *rmp4*) under the control of an inducible promoter, along with a vector control. Each clinical isolate is a unique combination of sequence type (ST) and capsule locus type (KL), and the strain set included ten STs and seventeen KL types (Table 2, Fig. S1). KPPR1S, a hypervirulent ST493 K2/O1 isolate with an *rmp3* locus, was used as a reference strain. HMV was assessed after *rmp* expression was induced (Figs. 1A, S2). In this assay, liquid cultures of HMV-negative strains easily sediment during low-speed centrifugation; HMV-positive cultures resist sedimentation.

**FIG 1.**
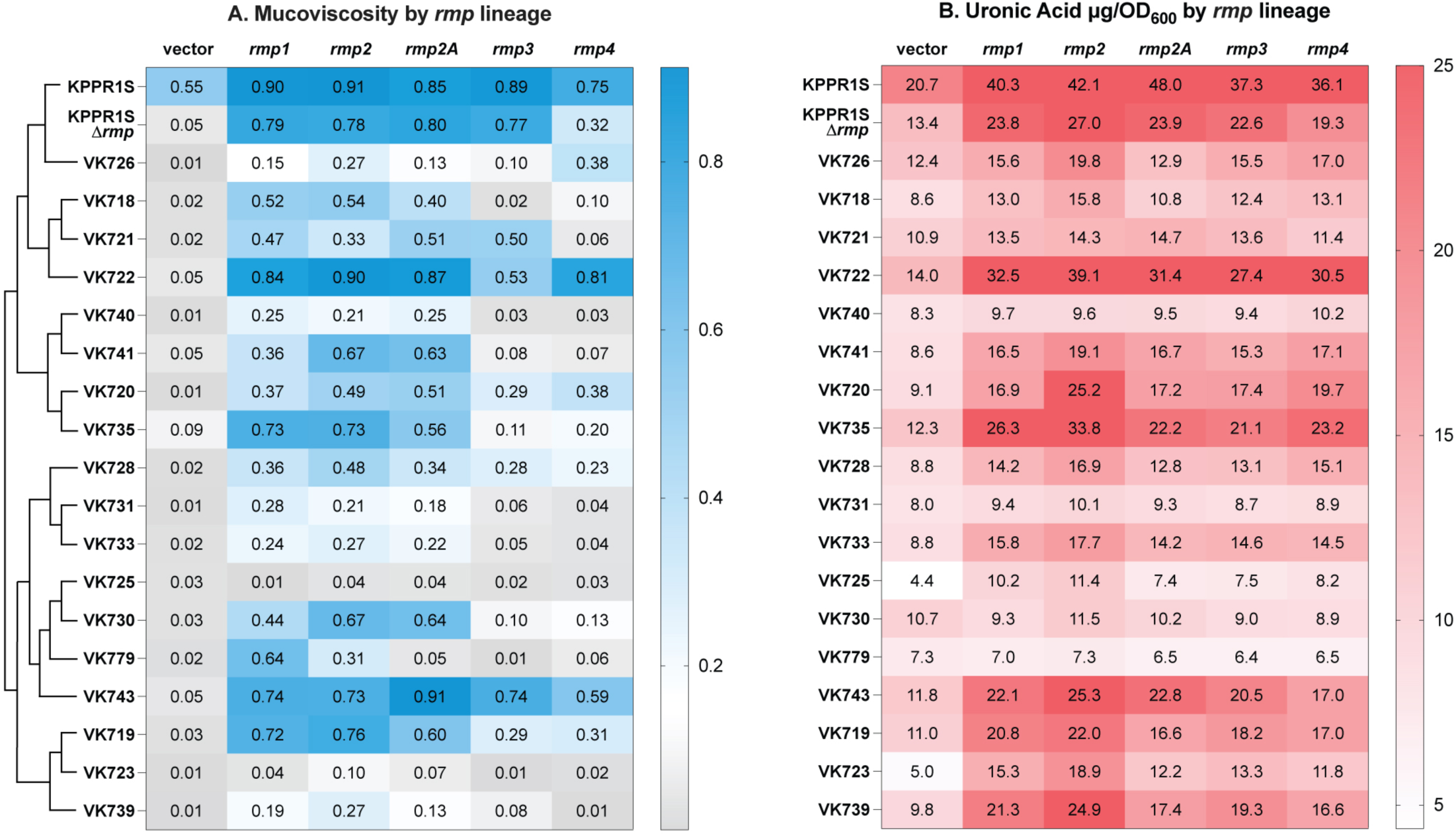
Hypermucoviscosity and capsule production vary by *rmp* lineage in classical strains. Strains were transformed with vector (pMWO-078) or lineage-specific pRmpADC. Strains were assayed for (A) sedimentation resistance (as post-spin/pre-spin OD_600_) and (B) uronic acid production (as µg/OD_600_) following 5.5 hour growth in LB with aTc to induce expression of plasmid-borne *rmp* genes as described in the Materials and Methods. The dendrogram in panel A illustrates evolutionary relationships between strains (the full tree is in Fig. S1). The dotted line (---) on the sedimentation resistance legend corresponds to 0.15, an established threshold for hypermucoviscosity [40]. Values are an average from three biological replicates. One-way ANOVA with Dunnett’s post-test was used to determine significance compared to vector controls; full dataset including statistics is available in Fig. S2.

**Table 1.** Plasmids used in this study.

| Plasmid | Relevant genotype | Reference or source |
| --- | --- | --- |
| pMWO-078 | Sp<sup>r</sup>; p15A ori cloning vector, *tetO* | [81] |
| pRmp1 / pKW215 | *rmpADC* from SGH10 in pMWO-078 | This work |
| pRmp2 / pKW216 | *rmpADC* from CIP 52.145 (B 5055, NTCC5055) in pMWO-078 | This work |
| pRmp3 / pKW186 | *rmpADC* from KPPR1S (ATCC 43816) in pMWO-078 | [82] |
| pRmp2A / pLPT059 | *rmpADC* from NCTC13669 in pMWO-078 | This work |
| pRmp4 / pKW222 | *rmpADC* from NCTC1936 in pMWO-078 | This work |
| pEW103 | Kan<sup>r</sup>; FRT flanks from pKD13 cloned into pUC18R6K-mini-Tn7T | [83] |
| pSMS013 | pEW103 with T1 terminator inserted at SacI site | This work |
| pSMS016 | pSMS013 with KPPR1S *rmp* (*rmp3*) | This work |
| pSMS026 | pSMS013 with SGH10 *rmp* (*rmp1*) | This work |
| pTNS2 | Ap<sup>r</sup>; R6K replicon; TnsABC+D expression vector | [84] |
| pFLP3 | Ap<sup>r</sup>; Tc<sup>r</sup>; Flp recombinase expression vector | [84] |

**Table 2.** Strains used in this study.

| Strain | Relevant genotype | ST | KL Type | Reference or source | Infection source |
| --- | --- | --- | --- | --- | --- |
| *E. coli* |  |  |  |  |  |
| DH5α | F<sup>-</sup> p80Δ*lacZ*M15 Δ(*lacZYA-argF*)U169 *deoP* *recA1* *endA1* *hsdR17* (r<sup>-</sup> m<sup>-</sup>) |  |  | Invitrogen |  |
| S17-1λpir | Tp<sup>r</sup> Str<sup>r</sup> *recA* *thi* *pro* *hsdR* *hsdM*<sup>+</sup> RP4::2-Tc::Mu::Km Tn7 λpir lysogen |  |  | [85] |  |
| *K. pneumoniae* |  |  |  |  |  |
| KPPR1S | ATCC 43816, Rif<sup>r</sup>, Str<sup>r</sup>; source of *rmp3* | ST493 | K2 | [86] |  |
| KPPR1SΔ*wcaJ* | KPPR1S, Δ*wcaJ* | ST493 | K2 | [27] |  |
| KPPR1SΔ*rmp3* | KPPR1S, Δ*rmpADC* | ST493 | K2 | [27] |  |
| KPPR1SΔ*rpmTn**rpm3* | Kan<sup>r</sup>; *rpm3* at *attTn7* site in KPPR1SΔ*rpm3* background | ST493 | K2 | This work |  |
| KPPR1SΔ*rpmTn**rpm3* Kan<sup>s</sup> | *rpm3* at *attTn7* site in KPPR1SΔ*rpm3* | ST493 | K2 | This work |  |
| VK718 | clinical isolate INF006 | ST35 | KL22 | [43] | other |
| VK719 | clinical isolate INF116 | ST36 | KL142 | [43] | urine |
| VK719*rpm1* | Kan<sup>r</sup>; *rpm1* at *attTn7* site | ST36 | KL142 | This work |  |
| VK719*rpm1* Kan<sup>s</sup> | *rpm1* at *attTn7* site | ST36 | KL142 | This work |  |
| VK719*rpm3* | Kan<sup>r</sup>; *rpm3* at *attTn7* site | ST36 | KL142 | This work |  |
| VK719*rpm3* Kan<sup>s</sup> | *rpm3* at *attTn7* site | ST36 | KL142 | This work |  |
| VK720 | clinical isolate INF119 | ST17 | KL38 | [43] | urine |
| VK720*rpm1* | Kan<sup>r</sup>; *rpm1* at *attTn7* site | ST17 | KL38 | This work |  |
| VK720*rpm1* Kan<sup>s</sup> | *rpm1* at *attTn7* site | ST17 | KL38 | This work |  |
| VK720*rpm3* | Kan<sup>r</sup>; *rpm3* at *attTn7* site | ST17 | KL38 | This work |  |
| VK720*rpm3* Kan<sup>s</sup> | *rpm3* at *attTn7* site | ST17 | KL38 | This work |  |
| VK721 | clinical isolate INF145 | ST35 | KL28 | [43] | disseminated |
| VK721*rpm1* | Kan<sup>r</sup>; *rpm1* at *attTn7* site | ST35 | KL28 | This work |  |
| VK721*rpm1* Kan<sup>s</sup> | *rpm1* at *attTn7* site | ST35 | KL28 | This work |  |
| VK721*rpm3* | Kan<sup>r</sup>; *rpm3* at *attTn7* site | ST35 | KL28 | This work |  |
| VK721*rpm3* Kan<sup>s</sup> | *rpm3* at *attTn7* site | ST35 | KL28 | This work |  |
| VK722 | clinical isolate INF168 | ST35 | KL16 | [43] | wound or tissue |
| VK723 | clinical isolate INF183 | ST36 | KL51 | [43] | urine |
| VK725 | clinical isolate INF201 | ST45 | KL24 | [43] | urine |
| VK726 | clinical isolate INF253 | ST101 | KL17 | [43] | urine |
| VK728 | clinical isolate INF346 | ST37 | KL12 | [43] | pneumonia |
| VK730 | clinical isolate KSB1_1D | ST45 | KL43 | [43] | rectal swab |
| VK731 | clinical isolate KSB1_6G | ST37 | KL153 | [43] | throat swab |
| VK733 | clinical isolate KSB1_7G | ST37 | KL15 | [43] | throat swab |
| VK733*rpm1* | Kan<sup>r</sup>; *rpm1* at *attTn7* site | ST37 | KL15 | This work |  |
| VK733*rpm3* | Kan<sup>r</sup>; *rpm3* at *attTn7* site | ST37 | KL15 | This work |  |
| VK735 | clinical isolate KSB1_9J | ST17 | KL25 | [43] | rectal swab |
| VK735*rpm1* | Kan<sup>r</sup>; *rpm1* at *attTn7* site | ST17 | KL25 | This work |  |
| VK735*rpm3* | Kan<sup>r</sup>; *rpm3* at *attTn7* site | ST17 | KL25 | This work |  |
| VK739 | clinical isolate KPN048 | ST307 | KL102 | This work | blood |
| VK740 | clinical isolate KPN071 | ST29 | KL24 | This work | urine |
| VK741 | clinical isolate KPN114 | ST29 | KL30 | [43] | urine |
| VK743 | clinical isolate KPN571 | ST147 | KL64 | This work | respiratory |
| VK779 | clinical isolate Kp209; KpnO2 | ST11 | KL110 | [87, 88] | urine |
| VK779*rpm1* | Kan<sup>r</sup>; *rpm1* at *attTn7* site | ST11 | KL110 | This work |  |
| VK779*rpm3* | Kan<sup>r</sup>; *rpm3* at *attTn7* site | ST11 | KL110 | This work |  |
| VK973 | clinical isolate BEI NR-55506 | ST37 | KL38 | [51] | urine |
| VK973*rpm1* | Kan<sup>r</sup>; *rpm1* at *attTn7* site | ST37 | KL38 | This work |  |
| VK985 | clinical isolate BEI NR-55518 | ST4270 | KL142 | [51] | urine |
| VK985*rpm1* | Kan<sup>r</sup>; *rpm3* at *attTn7* site | ST4270 | KL142 | This work |  |
| SGH10 / NCTC 14052 | source of *rpm1* | ST23 | K1 | [89] | liver abscess |
| Kp52.145 | source of *rpm2*. Derivative of B5055. | ST66 | K2 | [90, 91] | sputum |
| NCTC 13669 | source of *rpm2A* | ST3764 | KL1 | [37] | unknown |
| NCTC 1936 | source of *rpm4* | ST67<br>(subsp.<br>*rhinoscle*<br>*romatis*) | KL3 | [37] | unknown |

We previously observed that in KPPR1SΔ*rmp3, rmp* expression from the *rmp1*, *rmp2*, *rmp2A*, and *rmp3* lineages all conferred elevated HMV levels (included in Fig. 1, S2) [37]. However, expression of *rmp4* conferred a significantly lower level of HMV compared to the other lineages. We concluded that all *rmp* loci have some capacity to confer the HMV phenotype. When these same *rmp* plasmids were expressed in the cKp isolates, the changes in sedimentation resistance were much more varied (Figs. 1A, S2). Five strains became HMV positive in response to all *rmp* lineages, while eleven only responded to a subset of *rmp* lineages. Two strains did not exhibit HMV. Despite the variability, some patterns emerged. Expression of *rmp1* and *rmp2* converted more cKp strains to HMV-positive than the other lineages. In general, *rmp1* and *rmp2* conferred higher HMV levels than the other lineages, although this was not true for every isolate. Expression of *rmp3* and *rmp4* loci had less of an effect in the cKp strains than that of the other three lineages. Several isolates, such as VK718, VK733, VK739 and VK740, became HMV positive when *rmp1, rmp2,* and *rmp2A* were expressed, but not when *rmp3* or *rmp4* were expressed. Four strains became HMV positive from either *rmp3* or *rmp4* expression, but not both. While *rmp3* is the most divergent lineage of the five [37], the finding that both *rmp3* and *rmp4* converted fewer strains to HMV positive suggests that these patterns of mucoviscosity are not fully explained by the evolutionary relationships between *rmp* loci or between strains. These observations were validated by Pearson’s correlation analysis, which indicated a more positive correlation among HMV occurrence with *rmp1/2/2A* than *rmp3/4*. HMV occurrence with *rmp4* had only moderately positive correlation with all the other loci (Fig. S3A).

Previously published work demonstrated that HMV-negative strains produce diverse capsule chain lengths, as visualized by a smear on polyacrylamide capsule gels, whereas HMV-positive strains show more uniform capsule chain lengths [25, 39]. To examine capsule (CPS) properties of cKp expressing the *rmp* genes, we isolated CPS from the same cultures we had assessed for HMV and performed capsule gel electrophoresis. The HMV-positive classical transformants all showed distinct bands that indicate uniformly sized CPS chains (modal bands), whereas the HMV-negative transformants showed smears indicating varied chain lengths (Fig. S2). The presence of a modal band also corresponded to a sedimentation resistance at or above the threshold 0.15 that was established to distinguish HMV positive from HMV negative [40]. The size of the modal bands varied from strain to strain but were consistent within each classical strain background, regardless of *rmp* source or HMV level. Together, these sedimentation resistance and CPS profiling data show that both *rmp* lineage and strain genetic background play a role in HMV status

### Ectopic *rmp* expression confers hypercapsule independent of HMV

To determine if *rmp* expression in the cKp strains also resulted in hypercapsule, we assayed cKp transformants described above for capsule production by quantifying uronic acid (UA) (Figs. 1B, S1). We found that *rmp* expression from all lineages increased capsule production independent of HMV status. In fact, *rmp* expression conferred hypercapsule in most isolates, including two that did not become hypermucoviscous with any *rmp* locus (VK723 and VK725). Pearson’s correlation analysis showed no correlation between HMV and UA levels (Fig. S3A). Overall, these results are consistent with previous reports that HMV and capsule production are separable traits [27, 34, 41], and further show that both strain background and *rmp* lineage play a role in CPS phenotypes. As there did not appear to be any correlation between HMV and UA phenotypes within each strain or between which strains became HMV positive, we performed a principal component analysis (Fig. S3B). This showed the isolates were scattered over the plot area and validated that the data presented in Figs. 1 and S2 cannot be easily explained by evolutionary relatedness.

### Chromosomal *rmp* integration with native promoter validates role of *rmp* lineage

Having demonstrated that cKp isolates exhibit hv-associated phenotypes in response to the ectopic expression of *rmp*, we wanted to explore the effects of *rmp* acquisition on virulence. To circumvent the challenges of using inducible promoters and use of plasmid-borne genes *in vivo*, we opted to utilize the Tn7 system to introduce *rmp* operons with native promoters onto the chromosomes of a diverse subset of cKp isolates (Fig 2A). Two representative *rmp* loci were selected for integration based on their distinct effects on cKp isolates, the *rmp1* locus from SGH10 and the *rmp3* locus from KPPR1S. Both *rmp1* and *rmp3* loci have been detected in multiple clonal groups [42]. The *rmp1* lineage is particularly clinically relevant, as it is the most frequently found *rmp* lineage in hypervirulent isolates and is located on KpVP-1, the dominant *K. pneumoniae* virulence plasmid [43]. KpVP-1 has also been stably maintained in convergent strains of concern [37, 44, 45]. The *rmp3* lineage, which is found on the ICE*Kp1* element, is the most divergent from the other *rmp* lineages [37]. The recipient cKp isolates were selected because they all differentially responded to the ectopic expression of *rmp* loci; while *rmp1* expression conferred HMV and hypercapsule to every selected strain, the effects of *rmp3* expression were more variable. The chosen isolates include diverse sequence types and capsule loci, and were isolated from a variety of infection sources (Table 2, Table S2).

**FIG 2.**
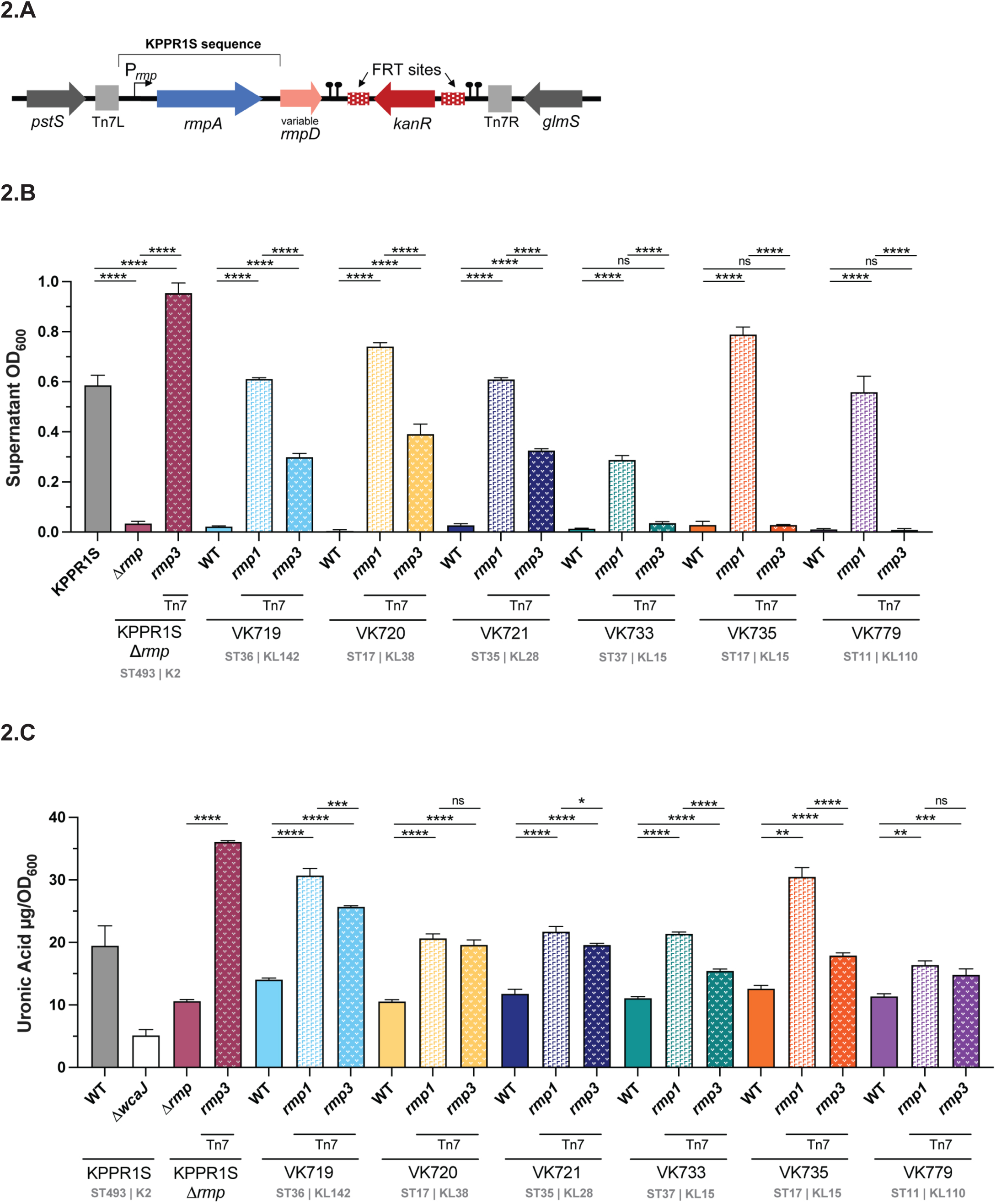
Chromosomal *rmp* integration affects HMV and capsule production in classical strains. (A) The *rmp1* or *rmp3* locus was integrated onto the chromosome at the *att*Tn*7* site and confirmed by PCR. After chromosomal *rmp* integration, strains were assayed for (B) mucoviscosity and (C) uronic acid production. *ΔwcaJ* is a capsule deficient derivative of KPPR1S included as a negative control. Data were obtained from at least three biological replicates following a 5.5 hour growth in LB as described in Methods. Bars represent average values while error bars indicate standard deviations. One-way ANOVA with Tukey’s post-test was used to determine significance compared to parental strains. *, *P* < 0.05; **, *P* < 0.01; ***, *P* < 0.001; ****, *P* < 0.0001; ns, not significant. Sequence (ST) and capsule (K or KL) type for each strain is listed.

The *rmp* integrants were evaluated for HMV and capsule production as described above (Figs. 2B, 2C). In general, single copy chromosomal integration of *rmp* loci in the cKp strains conferred the same pattern of HMV and UA phenotypes as ectopic expression of *rmp* loci shown in Fig. 1. Integration of *rmp1* converted all six isolates to HMV positive, while *rmp3* integration converted three isolates to HMV positive. Both lineages increased capsule production in all six strains. As a control, we integrated *rmp3* at the Tn7 site of the KPPR1SΔ*rmp3* mutant and found this strain showed higher levels of HMV and UA than its wildtype parent. This change in genetic location could impact expression due to differences in supercoiling or other *rmp* regulatory mechanisms. In general, the differential effects of *rmp1* and *rmp3* integration were less distinct in the capsule data compared to the HMV data. While *rmp1* integration conferred universally higher levels of HMV, *rmp1* and *rmp3* both increased capsule production to similar levels in most strains. Together, these data illustrate that *rmp* lineage can impact virulence-associated phenotypes in cKp.

### HMV correlates with hv-associated immune evasion phenotypes

HMV has been shown to reduce adherence to mammalian cells and to block phagocytosis by macrophages, which could contribute to immune evasion [27, 46]. To determine if *rmp* acquisition by cKp alters interactions with host cells we performed an adherence assay with the macrophage-like J774A.1 cell line for the panel of isolates with chromosomally integrated *rmp1* and *rmp3*. For each strain tested, adherence was inversely correlated with HMV level (Fig. 3). HMV-positive *rmp* integrants adhered at significantly lower levels than their HMV-negative parental strains. Adherence and internalization data typically display similar patterns, as blocking adherence and resisting phagocytosis are both HMV-dependent processes. We tested a subset of our cKp *rmp* integrants to determine if *rmp* integration impacted phagocytosis resistance. As expected, HMV-positive integrants were more resistant to internalization than their parental strains (Fig. S4).

**FIG 3.**
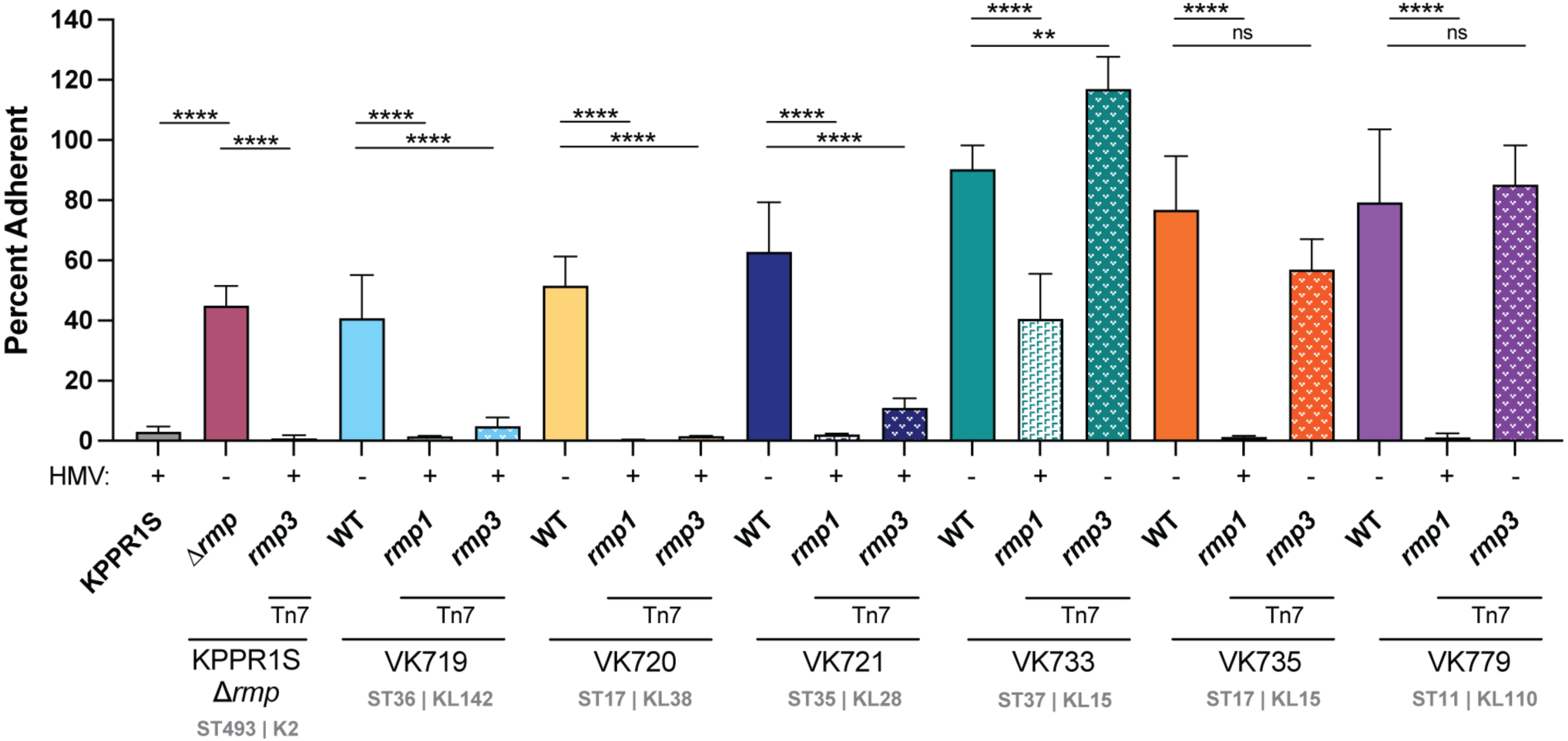
*rmp* acquisition blocks adherence in an HMV-dependent manner. Adherence of bacteria to J774A.1 cells treated with cytochalasin D to prevent phagocytosis was determined as described in the Materials and Methods. Data were obtained from at least three biological replicates. Bars represent average percent adherence, error bars indicate standard deviations. One-way ANOVA with Tukey’s post-test was used to determine significance compared to parental strains. ns, not significant; ***, *P* < 0.001; ****, *P* < 0.0001.

### HMV enhances virulence of some, but not all, classical strains

The acquisition of hypervirulence-associated genes and their associated phenotypes can potentially alter the virulence of classical strains, but it is unknown which genes are sufficient to increase virulence [44, 47–49]. Having established that *rmp* integration confers HMV and hypercapsule in many classical isolates, we wanted to investigate whether *rmp* acquisition increases virulence in a pneumonia model. Based on previous experiments with HMV-positive and HMV-negative strains, and the cell adherence data described above, we predicted that HMV-positive *rmp* integrants would have an increased ability to colonize mouse lungs and to disseminate to internal organs compared to their parental cKp strains. To test this, we inoculated mice intranasally with 2 x 10^7^ CFU of bacteria, then collected lungs and spleens at 24 and 72 hours post infection (hpi) to measure the bacterial burden.

In this pneumonia model, the parental clinical isolates all colonized similarly to each other at 24 hpi and bacterial burdens were appreciably lower by 72 hpi (Fig. 4). However, the colonization of HMV-positive *rmp* integrants varied substantially. We first tested VK719, VK720, and VK721 strains and their isogenic integrants, as these three cKp backgrounds are HMV positive with both *rmp1* and *rmp3* integration. This set of six HMV-positive strains all displayed high levels of sedimentation resistance and did not adhere to cells in culture, but only the VK719 *rmp* integrants caused disease in the pneumonia model (Fig. 4A). VK719*rmp1* and VK719*rmp3* both showed a significant increase in colonization in the lung compared to VK719 at 24 hpi. By 48 hpi, these strains had established severe infections that had spread to internal organs. The bacterial burden of the HMV-positive *rmp* integrants in the lung was approximately five logs higher than that of the VK719 parental strain. In contrast, the HMV-positive VK720 and VK721 integrants showed no change in colonization compared to their HMV-negative parental strains (Fig 4B, 4C).

**FIG 4.**
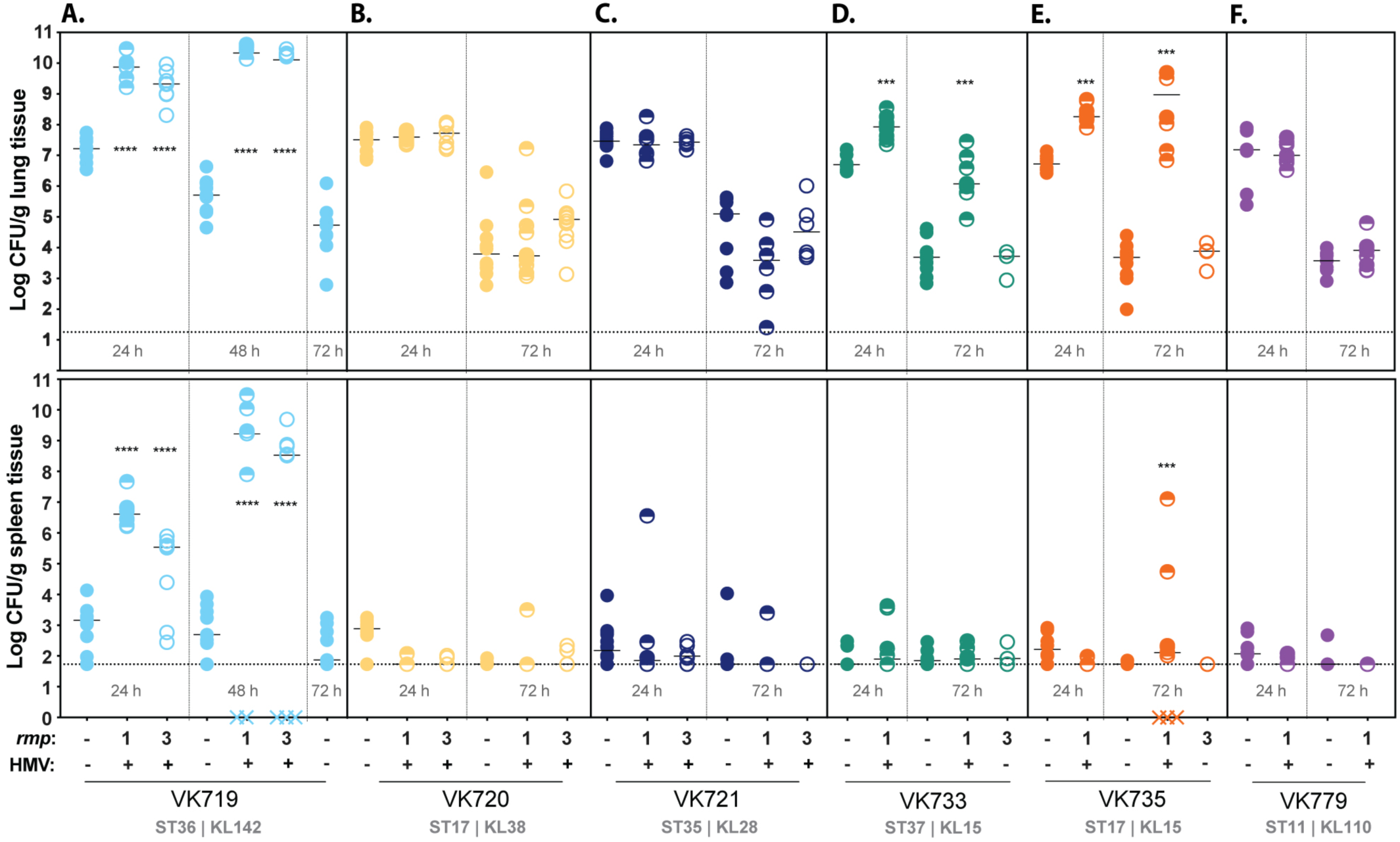
*rmp* integration and HMV conversion does not universally contribute to virulence. C57BL/6J mice were inoculated with 2 × 10^7^ CFU of the indicated strains; lungs (top) and spleens (bottom) were harvested at 24, 48, or 72 hpi for bacterial enumeration. Each symbol represents one mouse, solid lines indicate median values, and dotted lines represent the limit of detection. Symbols (X) on the X-axis represent mice that died prior to timepoint. HMV status is marked by - or + along the X-axis. HMV-negative cKp integrants were not tested at 24 h. Mann-Whitney test was applied to determine significance compared to parental strains. ***, *P* < 0.001; ****, *P* < 0.0001. Comparisons not indicated were not significant.

Having observed that HMV gain alone was not predictive of enhanced virulence, we expanded our colonization experiment to include additional integrants to determine if this observation was more generalizable. Three cKp isolates, VK733, VK735, and VK779, are HMV positive with *rmp1* integration, but not with *rmp3*. Given their limited capacity to become HMV positive, we were unsure how they would behave *in vivo*.

VK733*rmp1* and VK735*rmp1* both showed a significant increase in colonization compared to their parental strains at 24 and 72 hpi (Fig. 4D, 4E). Neither strain colonized as robustly as VK719*rmp1* or VK719*rmp3*, however, and while VK735*rmp1* spread to spleens, VK733*rmp1* did not. This suggests that *rmp* lineage can impact virulence through conferring HMV: HMV-positive VK733*rmp1* and VK735*rmp1* were virulent, while HMV-negative VK733*rmp3* and VK735*rmp3* were not. Finally, HMV-positive VK779*rmp1* was found to be avirulent (Fig. 4F). This further illustrates that HMV gain is not sufficient to increase virulence. In looking at available strain features (Table S2), we did not find anything that was exclusively unique among the strains that showed increased virulence versus those that did not. However, we did note that all three of the strains that gained HMV-dependent virulence (VK719, VK733, and VK735) contained the yersiniabactin-encoding genes (*ybt*), whereas only one of the three avirulent strains (VK720) contained *ybt* (Table S2).

Given that the majority of HMV-positive *rmp* integrants did not display an increase in virulence, we wanted to assess if *rmp* gene expression *in vivo* could be a determining factor. We infected mice with VK719*rmp3*, VK720*rmp3*, and VK721*rmp3*, as in Fig. 4 and collected lungs at 24 hpi for RNA extraction and qRT-PCR. Transcripts for *rmpA* were detected in lung samples from mice infected with these three strains, but we could not confidently compare expression between strains (Fig. S5). While not definitive, this result suggests that *rmp* is expressed *in vivo* in both virulent and avirulent strains, and that the colonization differences among these integrants are likely due to factors other than *rmp* expression. Taken together, these data show that the HMV phenotypes of these isolates *in vitro* cannot predict virulence *in vivo*.

### Capsule type does not predict *rmp-*dependent virulence

A recent study highlighted the importance of capsule type to *K. pneumoniae* virulence [50]. Given that HMV is dependent on capsule production, and our clinical isolates have highly diverse KL types, we decided to explore if capsule type might impact colonization phenotypes. We acquired additional clinical isolates with the same KL types as two of our original strains and integrated *rmp1* onto their chromosomes (VK985 and VK973) [51]. VK719 (ST36) and VK985 (ST4270) are KL142, VK720 (ST17) and VK973 (ST37) are KL38, and all four strains were isolated from urinary samples. When assayed for sedimentation resistance and capsule production, both VK985*rmp1* (KL142) and VK973*rmp1* (KL38) were HMV positive and produced more CPS than their parental strains (Fig. 5). However, in our pneumonia model, we found that VK985*rmp1* did not colonize lungs better than its parental strain, despite being the same KL type as one of the most virulent strains, VK719*rmp1* (Fig. 5C). In fact, at 24 hpi VK985*rmp1* was recovered from lungs at lower numbers than its parental strain. VK973*rmp1* was also avirulent but behaved similarly to its parental strain. Though limited, these results suggest that KL type alone cannot predict *rmp*-dependent increases in virulence. These data further point to the importance of strain background to any *rmp-*dependent virulence changes.

**FIG 5.**
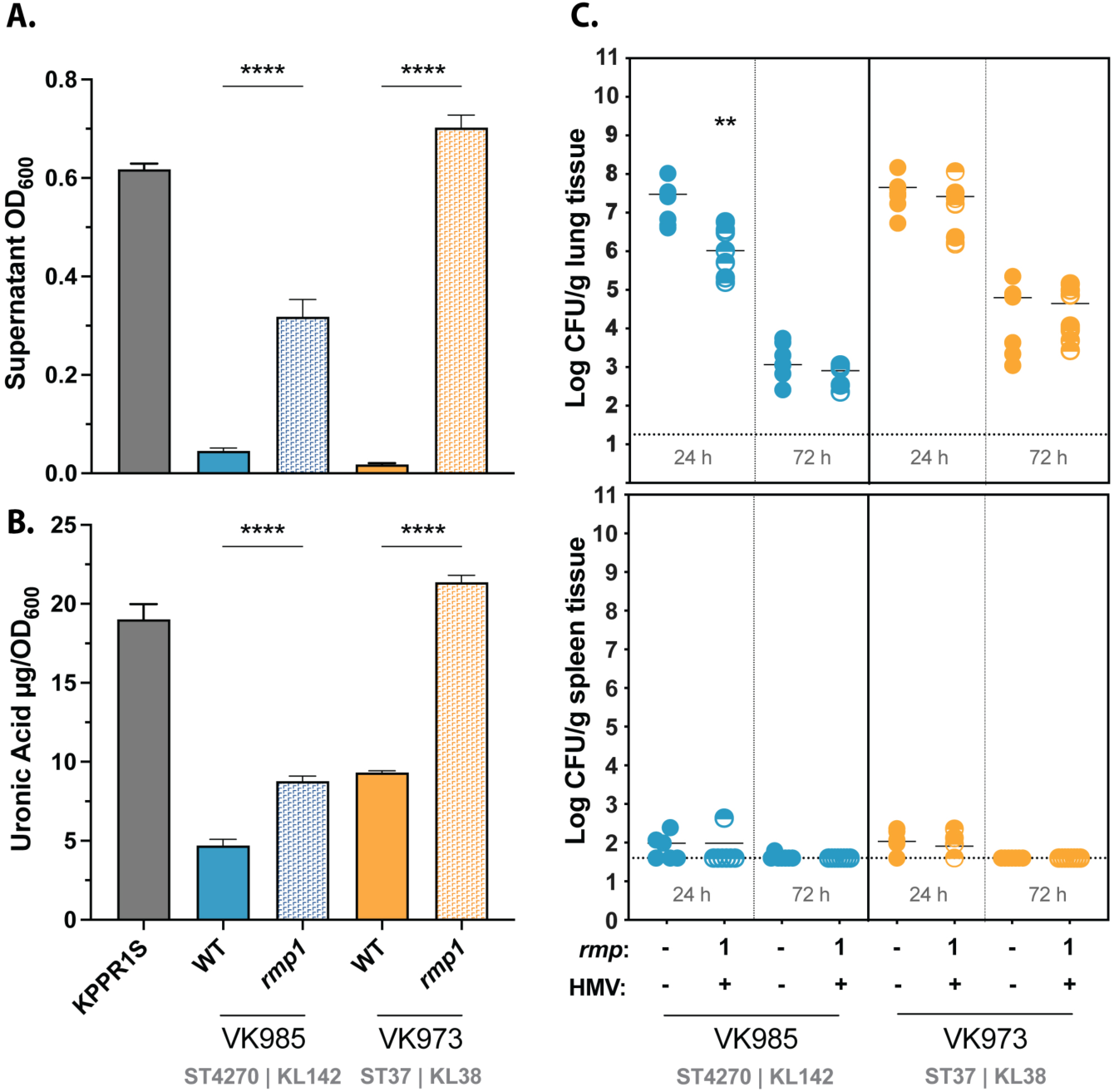
KL type alone does not determine *rmp-*dependent virulence of cKp isolates. Strains were assayed for (A) mucoviscosity, (B) uronic acid production, and (C) virulence in a mouse model. Mucoviscosity and uronic acid data were obtained from three biological replicates following growth for 5.5 hours in LB as described in Methods. Bars represent average values, error bars indicate standard deviations. One-way ANOVA with Tukey’s post-test was used to determine significance compared to parental strains. *, *P* < 0.05; **, *P* < 0.01; ***, *P* < 0.001; ****, *P* < 0.0001. C57BL/6J mice were inoculated with 2 × 10^7^ CFU of the indicated strains; lungs (top) and spleens (bottom) were harvested at 24 and 72 hpi for bacterial enumeration. Each symbol represents one mouse, solid lines indicate median values, and dotted lines represent the limit of detection. Mann-Whitney test was applied to determine significance compared to parental strains. **, *P* < 0.005. Comparisons not indicated were not significant.

## DISCUSSION

The growing threat of *Klebsiella pneumoniae* strains that are both highly virulent and antibiotic-resistant presents an urgent need for a better understanding of how hv-associated genes confer hypervirulent phenotypes [16, 52]. The presence of five genes (*rmpA*, *rmpA2*, *peg-344*, *iucA*, and *iroB*), along with the HMV phenotype, are typically used to define hvKp [53]. While these biomarkers are often used to identify convergent strains, they are not always predictive of virulence. In a recent study of convergent isolates, several strains with at least one of these hv-associated genes did not display hypervirulence in a mouse model, implying that acquisition of these genes was not sufficient for hypervirulence [49]. However, one convergent strain demonstrated virulence potential on par with hvKp strains, and that isolate was the only HMV-positive strain tested [49]. Given the virulence variability, it was recently proposed to only define strains as hvKp if they have all five biomarkers or have been shown to be hypervirulent in an appropriate infection model [54].

In studies on the hvKp strain KPPR1S, the *rmpADC* locus and HMV are both required for full virulence in a lung infection model [27, 55]. This is consistent with recent work demonstrating that *rmp* genes have a strong impact on *K. pneumoniae* lethality in mice [55]. In this present study, we aimed to determine if *rmpADC* acquisition and HMV gain alone were sufficient to confer hypervirulence to cKp. While other studies have characterized convergent isolates that have several hv-associated genes, we focused on the impact of *rmp-*dependent HMV. By introducing different *rmp* loci into a broad set of cKp isolates, we were able to draw general conclusions about how these genes can affect virulence-associated phenotypes, as well as actual virulence. After characterizing a large collection of cKp strains with *rmp* genes from different lineages, we found that HMV-positive phenotypes observed *in vitro* are not predictive of increased virulence in the mouse pneumonia model.

We first surveyed eighteen classical clinical isolates transformed with low-copy plasmids expressing representative *rmp* loci from each of the five *rmp* lineages. We found that ectopically expressing *rmp* genes in these isolates conferred a variety of HMV and hypercapsule phenotypes, consistent with a previous characterization of *rmp* function in a single cKp isolate [56]. While *rmp3* and *rmp4* converted the fewest strains to HMV positive and *rmp1* and *rmp2* converted the most, there was no consistent pattern. Some classical strains responded to all *rmp* lineages, some responded to none, and several isolates responded in entirely unique ways. All HMV-positive integrants displayed a known property of HMV: more uniform CPS chain lengths. However, the molecular weight of CPS chain lengths varied widely across cKp strains, and the size of the modal band had no clear correlation with sedimentation resistance, HMV capacity, or virulence potential. The interaction of RmpD with Wzc, a conserved capsule biosynthesis protein involved in capsule polymerization and export, plays a role in the change in capsule chain length associated with HMV [25]. The *wzc* gene is strikingly diverse between different capsule loci [57], and *rmpD* varies across *rmp* lineages [37]. Given the wide variety of K types in our clinical isolates, it is not surprising that these diverse RmpD-Wzc pairings might impact HMV phenotypes.

We observed that *rmp* expression increased capsule production independent of HMV. This validates previous observations that HMV and elevated capsule production are separable traits [34], and shows that RmpC can affect capsule production in strains with a broad array of capsule types. Elevated capsule production is an *rmpC-*dependent phenotype [34], and therefore suggests that all three *rmp* genes were expressed in these strains. This supports the idea that varied HMV phenotypes shown in Fig. 1A could be due, at least in part, to functional RmpD differences rather than variation in expression. Interestingly, there is less diversity across *rmpC* genes than across *rmpD* genes [37]. The higher level of sequence similarities might explain the broader functionality of RmpC across the *rmp* loci tested. Together, the HMV and capsule data show that *rmp* lineage can influence the characteristics of hv-associated phenotypes in cKp.

To facilitate virulence studies, we integrated *rmp1* or *rmp3* onto the chromosomes of a subset of cKp isolates. HMV and capsule assays of these integrants validated the importance of *rmp* lineage, as *rmp1* integration conferred higher HMV and UA levels in cKp isolates than *rmp3*. As observed with hvKp strains, HMV-positive cKp *rmp* integrants exhibited very low levels of adherence to mammalian cells, supporting previous data suggesting that HMV plays a role in immune evasion [27, 28]. When tested in a mouse pneumonia model, acquisition of *rmp* conferred a wide range of changes in virulence: VK719*rmp1* and VK719*rmp3* spread systemically and caused severe infections by 48 hpi, VK735*rmp1* spread systemically and caused severe infections by 72 hpi, and VK733*rmp1* established an infection, but did not spread to the spleen in the time frame tested. While the majority of *rmp* integrants showed colonization patterns identical to their parental strains, *rmp* acquisition, in the absence of other hv-associated genes, can confer a colonization benefit in the lung for some strains.

Although HMV did not strictly correlate with virulence in the pneumonia model, HMV proved to be a prerequisite for elevated colonization in this set of strains. None of the HMV-negative *rmp* integrants colonized mice any differently than their parental strain. This is a significant observation, because there are reports of strains that exhibit mucoidy phenotypes without carrying *rmp* [58, 59], and mutations in the capsule export gene *wzc* are known to independently confer HMV [39, 50, 60–64]. Our data suggest that any virulence changes resulting from those *wzc* mutations would be strain-dependent, as our results show that HMV does not necessarily enhance lung colonization. Determining the genetic requirements of HMV-dependent virulence will be essential to understanding which strain lineages pose the highest threat to human health.

Despite the phenotypic and genetic diversity of our cKp integrants, we have not identified traits that consistently correspond to *in vitro* or *in vivo* strain behavior. Sequence type (ST) does not appear to be predictive of either HMV conversion or virulence capacity. We characterized a pair of ST17 strains and a pair of ST35 strains, and both pairs showed varying levels of HMV and virulence capacity. We also tested an ST11 strain, VK779. ST11 is a sequence type associated with strain convergence, and ST11 strains maintaining KpVP-1 have been identified [44, 48, 65–67]. However, VK779 only became HMV positive with two of the five *rmp* lineages, and both the HMV-positive VK779*rmp1* strain and its HMV-negative parent colonized mouse lungs to similar bacterial burdens and were readily cleared.

One genetic element that might influence the virulence potential of HMV-positive cKp integrants is the presence of yersiniabactin-encoding genes (*ybt*). Four of the eight clinical strain backgrounds we tested in mice have *ybt* genes, though they carry different alleles (Table S2) [23]. While all three of the strain backgrounds that exhibited elevated virulence contained *ybt,* only one of our three avirulent classical strain backgrounds (VK720) contained *ybt* (Table S2). Siderophore production is associated with virulence for both classical and hypervirulent strains [1]. Approximately 40% of classical strains carry yersiniabactin genes, and genomic surveys have demonstrated that *ybt* loci are more often detected in isolates associated with invasive infections [23]. Hypervirulent strains are much more likely to carry *ybt*, as well as genes encoding the additional virulence-associated siderophores salmochelin and aerobactin [22]. In hvKp studies, yersiniabactin and aerobactin mutants are attenuated in murine models of infection [68, 69]. In the hypervirulent KPPR1 strain, yersiniabactin was specifically shown to contribute to bacterial replication in murine respiratory tracts, in part by evading the host innate immune protein lipocalin 2 (LCN2) [70–72] However, in a recent study of convergent strains, the presence of yersiniabactin and aerobactin-encoding genes in HMV-negative strains did not appreciably affect virulence in a murine pneumonia model [49]. Collectively, these studies may indicate that both yersiniabactin and HMV are necessary for lung colonization. While our results show that *ybt* or HMV are not sufficient for enhanced virulence in every background, they may indicate that *ybt* plays a critical role in lung colonization in combination with HMV. The potential dual importance of both HMV and siderophore production to *K. pneumoniae* virulence, and the relationship between iron and *rmp* [73, 74], implies that HMV and metal acquisition could be linked. This strain set offers an opportunity to explore how *ybt* might contribute to HMV-dependent virulence.

As specific capsule types are associated with hypervirulent and convergent strains, we tested if capsule type could affect virulence potential. In our limited sampling, capsule type was not predictive of virulence potential. When we tested *rmp* integrants of the same KL142 type, they had different virulence patterns. It is important to note, however, that both isolates tested in Figure 5, VK985 (KL142) and VK973 (KL38), lack *ybt* genes, but their KL pairs tested in Figure 4 do contain *ybt* (Table S2)[51]. A more expansive study, including isogenic capsule swaps, will be needed to better understand how capsule type might influence HMV-dependent virulence capacity. A recent study demonstrated that hypervirulent strains become attenuated when the K1 or K2 *cps* locus is replaced with a K3 locus, highlighting the role of capsule type in determining virulence capacity [50]. The majority of identified hypervirulent strains have either a K1 or K2 capsule, and convergent strains also seem to be restricted to a limited number of capsule types. Reports of MDR-hvKp have included strains with K1, K2, K5, K16, K19, K20, K24, K47, K51, K54, K57, K62, K64, and KL108 capsules [10, 75]. Interestingly, one of the strains we tested that responded strongly to ectopic *rmp* expression was VK743, a KL64 strain (Figs. 1A, S1). While its native antibiotic resistances prevented further characterization, these data suggest that naturally occurring convergent strains may give us the best clues as to which genetic backgrounds are most primed for hypervirulence.

Overall, this study enhances our understanding of strain convergence and points to *K. pneumoniae* factors that, when combined with hypermucoviscosity, affect lung colonization. Given the strain diversity and phenotypic variability of our strain set, further examination of their genomes could reveal other determinants of lung colonization, and characterizing the role of *ybt* in both virulent and avirulent integrants might establish whether yersiniabactin promotes HMV-dependent virulence. Expanding the strain set to include Tn7 integrants with different combinations of *rmp* genes could also reveal how *rmpD* and *rmpC* separately contribute to virulence. Though this study provides insight into how classical strains can gain virulence, we cannot determine if these strain backgrounds would maintain virulence-associated mobile genetic elements in a natural environment. While we have demonstrated that strain background contributes to the effects of *rmp* expression, strain background also likely contributes to the dynamics of *rmp* maintenance and the acquisition of mobile virulence elements, as capsule type has been shown to influence plasmid conjugation [76]. More expansive studies are required to explore these questions due to the high genetic diversity of *Klebsiella* species.Further understanding the impact of the *rmp* genes on diverse genetic backgrounds is critical to addressing the threat of multi-drug resistant cKp strains that acquire virulence traits.

## MATERIALS AND METHODS

### Bacterial isolates

A diverse panel of cKp isolates was assembled from the *Klebsiella* Acquisition Surveillance Project at Alfred Health in Australia to include common clinical multidrug resistant (MDR) strain backgrounds (sequence types, STs) with a variety of capsule loci. Some capsule loci were specifically chosen for their association with emerging hypervirulence, and others because they are common human isolates but not MDR. The strains in this study all exhibited sensitivity to spectinomycin (to facilitate selection of transformants with pRmpADC), and the isolates chosen for virulence studies were also sensitive to kanamycin (to facilitate selection of Tn7*rmp* integrants). All isolates and relevant information are listed in Tables 2 and S2.

### Bacterial growth conditions

All strains were grown aerobically at 37°C in Luria-Bertani (LB) medium (10 g tryptone, 5 g yeast extract, and 10 g NaCl per liter). Kanamycin at 50 μg/ml (Kan_50_), tetracycline at 10 μg/ml (Tet_10_), or spectinomycin at 50 μg/ml (Spec_50_) were added to media as appropriate. For the ectopic *rmp* expression assays using pMWO-078-based plasmids, 100 ng/ml anhydrous tetracycline (aTc) was added to the medium at the time of subculture.

### Plasmid and strain construction

Plasmids and strains used in this study are described in Table 1 and Table 2; primers used for cloning and PCR are listed in Table S1. Expression plasmids were constructed by cloning *rmp* loci amplified from various *K. pneumoniae* strains into the SalI-BamHI sites of pMWO-078 using Gibson assembly (NEB). They were transformed into DH5α, confirmed by sequencing, and electroporated into *K. pneumoniae* strains as previously described [77]. Tn7 integrants were constructed by integrating *rmp* loci at the *att*Tn7 site of *K. pneumoniae* strains [78]. Briefly, *rmp* loci of interest were amplified and cloned into the XhoI-StuI sites of pSMS013, a pUC18Tmini-Tn7 derivative, by Gibson assembly (NEB). These plasmids were electroporated into *K. pneumoniae* strains along with pTNS2 as previously described, recovered overnight at 37°C, and plated on Kan_50_ plates [77]. Chromosomal *rmp* insertion was confirmed by colony PCR. In selected strains, the Kan cassette was removed using pFLP3, as previously described [78].

### Assessment of HMV and capsule production

Mucoviscosity of liquid cultures was determined using a sedimentation assay as previously described [26, 34]. Overnight cultures were diluted to an OD_600_ of 0.2 and grown for 5.5 h at 37°C. Cultures were normalized to 1 OD_600_/ml and centrifuged at 1,000 × *g* for 5 min. The OD_600_ values of the post-spin supernatants were divided by the pre-spin OD600 values to determine sedimentation resistance. Capsule production was determined by quantifying uronic acid as described previously [79]. Cultures were grown as for the mucoviscosity assay. CPS was extracted from 500 μl of culture with 1% zwittergent, precipitated with 100% ethanol at 4°C overnight. CPS pellets were resuspended in tetraborate/sulfuric acid and boiled. 3-Phenylphenol was added and absorbance was measured at 520 nm. UA amounts were determined from a standard curve generated with glucuronolactone.

### Capsule SDS-PAGE

Capsule was visualized essentially as previously described [25]. Briefly, 1 OD of bacterial culture was collected after 5.5 hrs of subculture, an equal volume of 2x SDS-PAGE loading buffer was added and then boiled. Proteinase K (0.5 µg) was added and incubated overnight at 55°C. SDS-PAGE was performed with 10% acrylamide gels, followed by staining first with alcian blue then with silver (Pierce silver stain kit; Thermo Scientific) [80].

### Phylogenetic tree construction and correlation analyses

The genomes for each isolate were uploaded to Pathogenwatch (https://pathogen.watch/) and aligned against the *K. pneumoniae* reference core library (2,172,367 bp) to generate a matrix of pairwise distances and corresponding neighbor-joining tree. Pearson’s correlation coefficients (two-tailed) and principal component analysis were performed using Graphpad Prism v. 10 with the HMV and UA data for each strain in Figs. 1/S2.

### Adherence and internalization assays with J774A.1 cells

Adherence and internalization assays were performed as previously described [27, 77]. Bacteria were added to J774A.1 at an MOI of 50 and incubated for 1 h. The cells were then washed with PBS and treated with 0.5% Saponin. Recovered CFU were compared to the inoculum CFU. For adherence assays, the J774A.1 cells were pretreated with cytochalasin D for 1 h prior to inoculation to prevent phagocytosis. For internalization assays, the wells were treated with 200 μg/ml gentamicin for 30 min following the 1-h coincubation, then J774A.1 cells were lysed and bacterial CFUs were plated as for the adherence assay.

### Murine pneumonia model

All animal experiments were approved by the Institutional Care and Use of Committee of UNC-Chapel Hill and were conducted in accordance with the Guide for the Care and Use of Laboratory Animals of the National Institutes of Health. C57BL/6J mice (Jackson Laboratories, Bar Harbor, ME) were anesthetized with 200 µl of a 0.8 mg/ml ketamine and 1.3 mg/ml xylazine solution and inoculated with 2 × 10^7^ CFU intranasally as described previously [77]. Mice were sacrificed at 24, 48, or 72 hpi by intraperitoneal injection of 200 µl sodium pentobarbital (52 mg/ml). Lungs and spleens were removed, weighed, homogenized, and plated to determine CFU/gram of tissue.

### qRT-PCR from *in vivo* RNA

Mice were inoculated with 2 × 10^8^ CFU bacteria intranasally and sacrificed at 24 h. Lungs were removed and each of the large lobes were placed in 1 mL RNA Later (Thermo Fisher Scientific) and stored at -80°C. To extract RNA, tissues were transferred to tubes containing PBS and 5 silica beads and homogenized at 5,600 rpm for a total of 3 minutes in a Precellys 24 homogenizer (Bertin Technologies). Homogenates were then passed through a 40 µm cell strainer (VWR) into a 50 mL conical. This filtrate was then passed through a 5 µm syringe filter (Sartorius). Total RNA was extracted from the filtrate with Trizol reagent (Invitrogen) following the manufacturer’s instructions, with an additional bead beating step, as previously described [77]. Contaminating DNA was removed with DNA-*free* Turbo, and cDNA was synthesized using an iScript cDNA synthesis kit (Bio-Rad). qRT-PCR was performed using SsoAdvance SYBR green Supermix (Bio-Rad) in a CFX96 RealTime system (Bio-Rad). The transcript levels for the *rmpA* target gene were compared to those from the *gyrB* gene.

### Statistics and replicates

Statistical tests for each dataset were performed by GraphPad Prism 10 and are provided in the figure legends. A minimum of two assays were performed for every experiment, each with at least three biological replicates. Representative experiments are presented for the *in vitro* assays; combined experiments are presented for the *in vivo* assays.

## ACKNOWLEDGMENTS

We thank Kelly Wyres for guidance and input, Sarah Rowe Conlon and Ammar Zafar for insightful discussions and manuscript feedback, Sylvain Brisse for permission to work with strain 52.145, and Suzan H. M. Rooijakkers for providing VK779 (Kp209). Thank you to Logan Treat, Alex Tufano, and members of Rita Tamayo’s lab for their support. This work was supported by R01 AI148197 to V.L.M. from NIAID. This material is partially based upon work supported by the National Science Foundation Graduate Research Fellowship Program under Grant No. DGE-2439854 to L.A.K. Any opinions, findings, and conclusions or recommendations expressed in this material are those of the author(s) and do not necessarily reflect the views of the National Science Foundation.

## AUTHOR CONTRIBUTIONS

Conceptualization of study: KAW, VLM, SMS, MMCL and KEH

Data acquisition: SMS, TAM, LAK, VLM, MMCL and KAW

Analysis and interpretation of data: SMS, KAW, VLM, MMCL and KEH

Initial drafting of manuscript: SMS with input from VLM and KAW

All authors edited and approved the submitted manuscript.

---

**Table S1.**
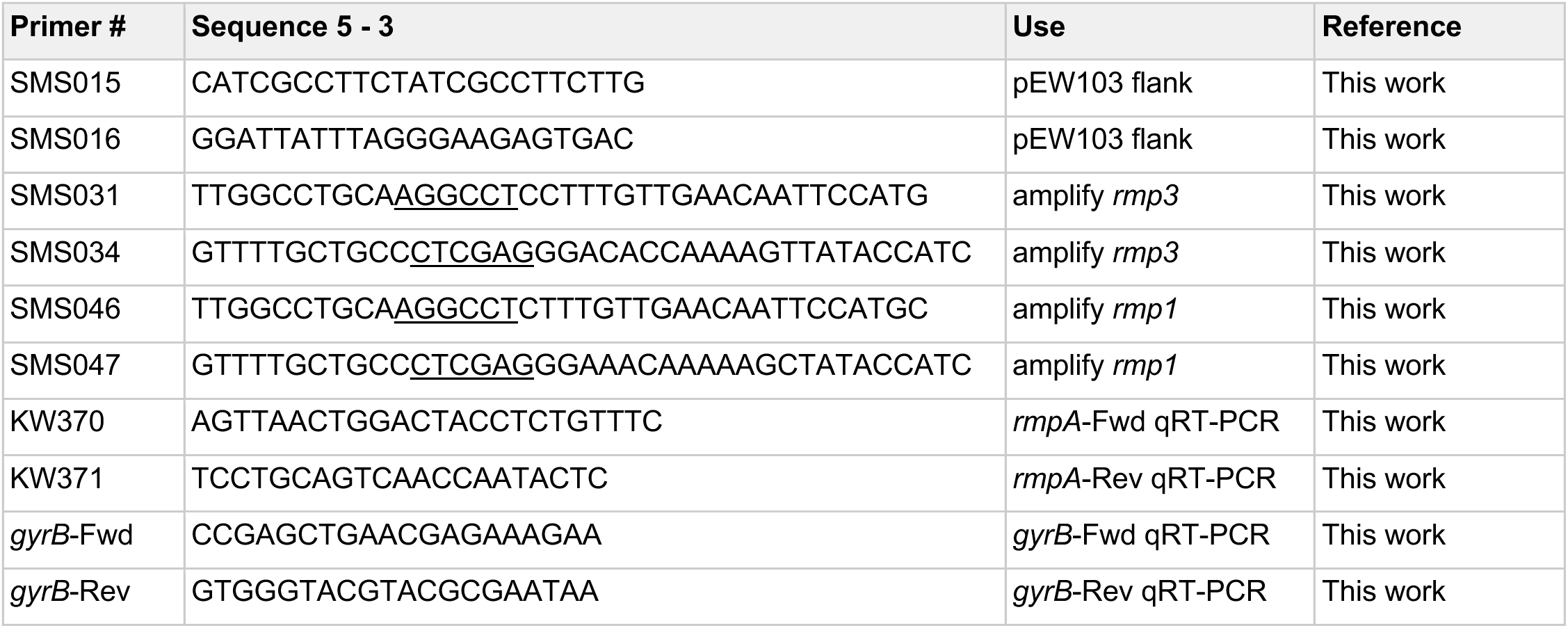
Primers used in this study.

